# Inferring population size history from large samples of genome wide molecular data - an approximate Bayesian computation approach

**DOI:** 10.1101/036178

**Authors:** Simon Boitard, Willy Rodríguez, Flora Jay, Stefano Mona, Frédéric Austerlitz

## Abstract

Inferring the ancestral dynamics of effective population size is a long-standing question in population genetics, which can now be tackled much more accurately thanks to the massive genomic data available in many species. Several promising methods that take advantage of whole-genome sequences have been recently developed in this context. However, they can only be applied to rather small samples, which limits their ability to estimate recent population size history. Besides, they can be very sensitive to sequencing or phasing errors. Here we introduce a new approximate Bayesian computation approach named PopSizeABC that allows estimating the evolution of the effective population size through time, using a large sample of complete genomes. This sample is summarized using the folded allele frequency spectrum and the average zygotic linkage disequilibrium at different bins of physical distance, two classes of statistics that are widely used in population genetics and can be easily computed from unphased and unpolarized SNP data. Our approach provides accurate estimations of past population sizes, from the very first generations before present back to the expected time to the most recent common ancestor of the sample, as shown by simulations under a wide range of demographic scenarios. When applied to samples of 15 or 25 complete genomes in four cattle breeds (Angus, Fleckvieh, Holstein and Jersey), PopSizeABC revealed a series of population declines, related to historical events such as domestication or modern breed creation. We further highlight that our approach is robust to sequencing errors, provided summary statistics are computed from SNPs with common alleles.

## Introduction

Reconstructing the ancestral dynamics of effective population size is important in several contexts. From a long term evolutionary perspective, the history of population size changes can be related to various climatic or geological events, and reconstructing this history allows studying the impact of such events on natural species [1]. This demographic history also provides a statistical null model of neutral evolution that can subsequently be used for detecting loci under selection [2,3]. In conservation biology, the recent dynamics of effective population size in endangered species, as reconstructed from genetic data, can efficiently be used to decipher the time frame of a population decline, hence allowing to separate anthropogenic from natural factors [4].

Until recently, methods allowing to infer the history of population size changes from genetic data were designed for data sets consisting of a limited number of independent markers or non recombining DNA sequences [5-8]. However, the spectacular progress of genotyping and sequencing technologies during the last decade has enabled the production of high density genome-wide data in many species. New statistical methods accounting for recombination and scalable to the analysis of whole genome sequences are thus needed, in order to take advantage of this very rich source of information.

In this context, several promising approaches allowing to infer complex histories, including several tens of stepwise population size changes, have recently been proposed [9-13]. Some of them, called PSMC [9], MSMC [10] and diCal [11], are based on the Sequentially Markovian Coalescent (SMC or SMC’) models [14,15], an approximation of the classical coalescent with recombination [16], where coalescent trees are assumed to be Markovian along the genome. Thanks to this Markovian assumption, maximum likelihood estimates of past population sizes can be efficiently obtained from the observation of one (for PSMC) or several (for MSMC and diCal) diploid genomes. Another approach [12] is based on the length of Identity By State (IBS) segments shared between two chromosomes along the genome. Using an iterative search, it aims at finding a history of past population size changes for which the expected distribution of IBS segment lengths matches that observed in one diploid genome.

While the above methods take advantage of whole-genome data, they are so far restricted to the analysis of small sample sizes. In the case of SMC based methods, this implies a limited resolution for the estimation of recent population sizes. Indeed, the most recent time at which these methods can infer population size is determined by the time to the most recent coalescence event occurring in the sample, which is older for small samples. For instance in humans, PSMC cannot estimate population sizes more recently than 400 generations (10,000 years) before present (BP), and MSMC cannot estimate these sizes more recently than 40 generations (1,000 years) BP. The most recent time for which an inference is possible will differ between species. In populations with small recent population sizes, coalescence events will occur at a higher rate than in larger populations, so the inference of recent history will be more accurate. Inference approaches based on the distribution of IBS segment length may be less affected by the use of small samples. Using this approach, estimations of population size in the Holstein cattle breed were obtained from a single genome even for the first few generations BP [12], and were in good agreement with estimations obtained from pedigree information in this breed [17-19]. However, the accuracy of the IBS approach used in this study has not been formally validated using simulations.

Another concern of the above methods is their sensitivity to sequencing errors. False positive SNPs can lead to a strong overestimation of population sizes in the recent past, i.e. in the first few hundred generations BP, both with PSMC [9,12] and with the distribution of IBS segment length [12]. In contrast, false negative SNPs lead to underestimate population size at all time scales, but the magnitude of this effect is much weaker [12]. Efficient strategies for estimating these error rates and correcting the data accordingly have been proposed in [12]. However, the estimation step typically requires other sources of information than the sampled sequences, such as independent SNP chip data for the same individuals, which in many cases are not available. Phasing errors may also be an issue when inference is based on phased haplotype data, which is typically the case for MSMC [10] or diCal [11]. MSMC inference can also be based on unphased data, but this reduces the estimation accuracy [10].

Here we introduce a new statistical method named PopSizeABC, allowing estimating population size history from a sample of whole-genome sequences. One of the main motivations for developing this method is to take advantage of large sample sizes in order to reconstruct the recent history as well. Since statistical approaches based on the full likelihood of such samples seem currently out of reach, even with approximated models such as the SMC, we followed an Approximate Bayesian Computation (ABC) [20] approach, which simplifies the problem in two ways. First, this approach does not focus on the full likelihood of sampled genomes, but on the likelihood of a small set of summary statistics computed from this sample. Second, population size histories that are consistent with these observed summary statistics are inferred by intensive simulations rather than by complex (and generally intractable) mathematical derivations.

ABC is a popular approach in population genetics, which has already been applied to the analysis of large-scale population genetic data sets [21-25]. However, none of these previous studies tried to estimate complex population size histories involving a large number of population size changes. To address this question, we considered two classes of summary statistics: the folded allele frequency spectrum (AFS) and the average linkage disequilibrium (LD) at different physical distances. These two classes of statistics are very informative about past population size, and each of them is the basis of several inference approaches in population genetics [13, 26-31]. Therefore, combining them within an ABC framework seems very promising.

Applying our ABC approach to samples of 25 diploid genomes, simulated under a large number of random population size histories, we show that it provides, on average, accurate estimations of population sizes from the first few generations BP back to the expected time to the most recent common ancestor (TMRCA) of the sample. This result is confirmed by the study of several specific demographic scenarios, where our method is generally able to reconstruct the population size history from present time back to the expected TMRCA, while PSMC or MSMC reconstruct it only for a limited time window.

We then apply this method to samples of 15 or 25 genomes in four different cattle breeds, which reveals interesting aspects of cattle history, from domestication to modern breed creation. Through this application to a real data set, we also illustrate how sequencing and phasing errors, if not taken into account, can have a dramatic influence on the estimated past population sizes. Our method is actually insensitive to phasing errors, because it uses unphased data. In addition, we show that a simple modification in the choice of summary statistics makes it robust to sequencing errors.

## Results

### Overview of the approximate Bayesian computation (ABC) estimation procedure

Following several recent studies [9-12], we modeled population size history as a stepwise constant process with a fixed number of time windows, where population size was constant within each window but was allowed to change from one window to the next. Time windows were defined in generations, for instance the most recent window went from one to ten generations before present (BP), and the most ancient window started 130,000 generations BP. This model allows approximating all simple demographic scenarios generally considered in population genetics studies (constant size, linear or exponential growth or decline, bottleneck …), as well as a large range of more complex demographic scenarios, provided population size changes occurred more recently than 130,000 generations BP.

Our estimation procedure was based on the observation of n diploid genomes sampled from the same population. We summarized this data set using two classes of summary statistics: (i) the folded allele frequency spectrum (AFS) of the sample, which includes the overall proportion of polymorphic sites in the genome and the relative proportion of those polymorphic sites with i copies of the minor allele, for all values of i between 1 and *n*, and (ii) the average linkage disequilibrium (LD) for 18 bins of physical distance between SNPs, from approximately 500 bp to 1.5Mb. We generated a very large number of population size histories, by drawing the population size in each time window from a prior distribution. For each history, we simulated a sample of *n* diploid genomes and computed a distance between the summary statistics obtained from this simulated sample and those obtained from the observed sample. A given proportion (called tolerance) of the most likely histories was accepted based on this distance. Finally, the joint posterior distribution of population sizes was estimated from the population sizes of accepted histories. Different statistical approaches were compared for this last estimation step.

A detailed description of the model and of the ABC procedure described above is provided in the Methods.

### Accuracy of ABC estimation and relative importance of summary statistics

In order to optimize our ABC estimation procedure and to evaluate its average performance, we first applied it to a large number of genomic samples simulated under random population size histories. These pseudo-observed datasets (PODs) included 25 diploid genomes and 100 independent 2Mb-long regions. For each POD, population sizes were estimated by ABC, using 450.0 simulated datasets of the same size. These estimated values were compared with their true values for different tolerance rates and different ABC adjustment approaches to process the accepted histories. We found that the best procedure was to accept simulated histories with a tolerance rate of 0.005, to adjust their parameter values using a non linear neural network regression [32], and to summarize the resulting posterior distribution by its median. Indeed, point estimations of population sizes obtained by this procedure showed very small bias and the lowest prediction errors (PE) (Fig. 1). Moreover, the posterior distributions of population sizes in each time window were correctly estimated, as shown by the accuracy of the 90% credible interval (S1 Fig, left), while the size of this credible interval was much lower than that obtained by the other adjustment approaches considered (S1 Fig, right). We used this procedure throughout the remaining of this study.

**Figure 1.**
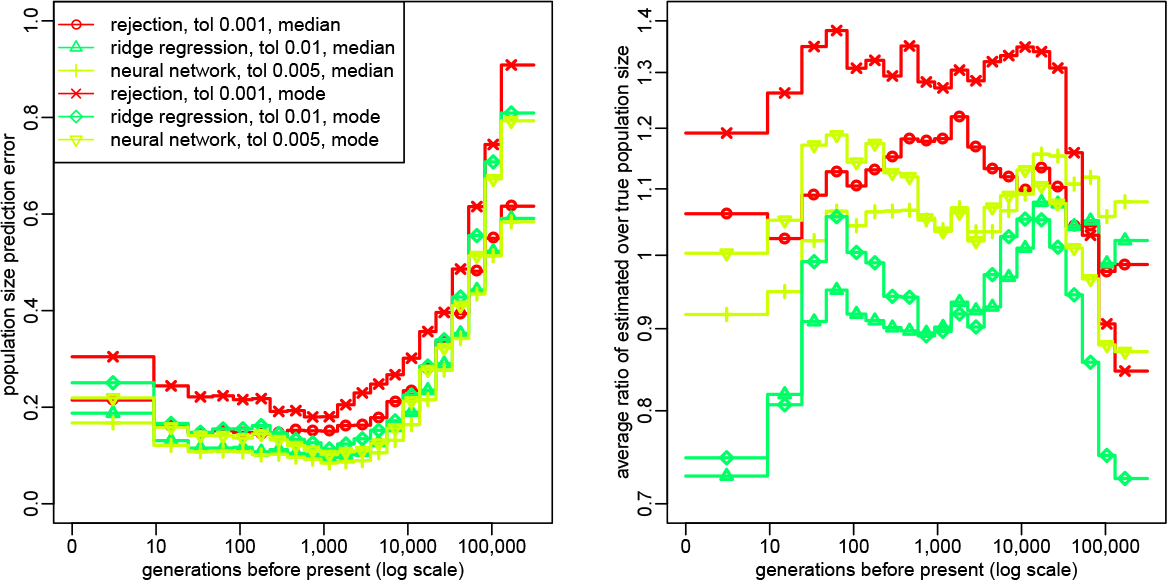
Optimization of ABC procedure: Prediction error (left panel) and bias (right panel) for the estimated population size in each time window, evaluated from 2,000 random population size histories (see Methods). Summary statistics considered in the ABC analysis were (i) the AFS and (ii) the average zygotic LD for several distance bins. These statistics were computed from *n* = 25 diploid individuals, using all SNPs for AFS statistics and SNPs with a MAF above 20% for LD statistics. The posterior distribution of each parameter was obtained by rejection, ridge regression [33] or neural network regression [32]. The tolerance rate used for each of these approaches was the one providing the lowest prediction errors, for different values from 0.001 to 0.05. Population size point estimates were obtained from the median or the mode of the posterior distribution. The prediction errors were scaled in order that point estimates obtained from the prior distribution would result in a prediction error of 1.

ABC provided accurate estimations of population sizes for a large range of times in the past (Fig. 1). The best results were obtained from 10 to 5.000 generations BP, where the prediction error was below 0.1: this means that the average distance between true and estimated population sizes for this period of time was more than 10 times smaller than if the population sizes were estimated from the prior distribution. In the very recent past (from 0 to 10 generations BP), this prediction error was slightly larger but remained below 0.2. The prediction error also increased for times more ancient than 5,000 generations BP, while remaining quite low (PE ≤0.3) until approximately 20,000 generations BP. This increase in prediction error above 5,000 generations BP can be related to a coalescence argument. At this time, the observed samples have coalesced to their common ancestor at most of the genomic regions, so the influence of demography on the current sample is reduced. Indeed, when rescaling time from generations to coalescent units (as described in Methods), we observed that the prediction error averaged over PODs started to increase shortly after the expected TMRCA (S2 Fig).

Our simulation study also highlighted the contribution of the different summary statistics. First, we found that population size history can be estimated quite well using either the AFS statistics alone or the LD statistics alone, but that combining the two classes of statistics clearly leads to the lowest PE for all time windows (Fig. 2, left). As some demographic histories were more difficult to estimate than others, the PE differed between histories, but we observed that combining AFS and LD statistics allowed to reduce these differences (Fig. 2, right). It also led to a reduction of the width of the 90% credible interval, as compared with the interval obtained using either class of statistics alone (S3 Fig). Another important advantage of combining AFS and LD statistics is to enable estimating the per site recombination rate. Indeed, the PE of this parameter was equal to 0.2 when using all statistics, versus 0.96 and 0.75 when using, respectively, AFS or LD statistics alone. Second, we found that using the polymorphic site AFS, i.e. the AFS without the overall proportion of SNPs, resulted in much higher PEs than using the full AFS (Fig. 2). Third, we observed that computing LD at each SNP pair as a correlation between two vectors of *n* genotypes or as a correlation between two vectors of 2*n* alleles was equivalent in terms of PE (S4 Fig). This result implies that, with our approach, using unphased data rather than phased data will not decrease the estimation accuracy. Besides, computing LD from SNPs with relatively frequent alleles (MAF ≥ 5-20 %) resulted in lower PEs than computing it from all SNPs (S4 Fig). In the following, LD statistics were always computed from genotype data at SNPs with a MAF above 20%, unless otherwise specified.

**Figure 2.**
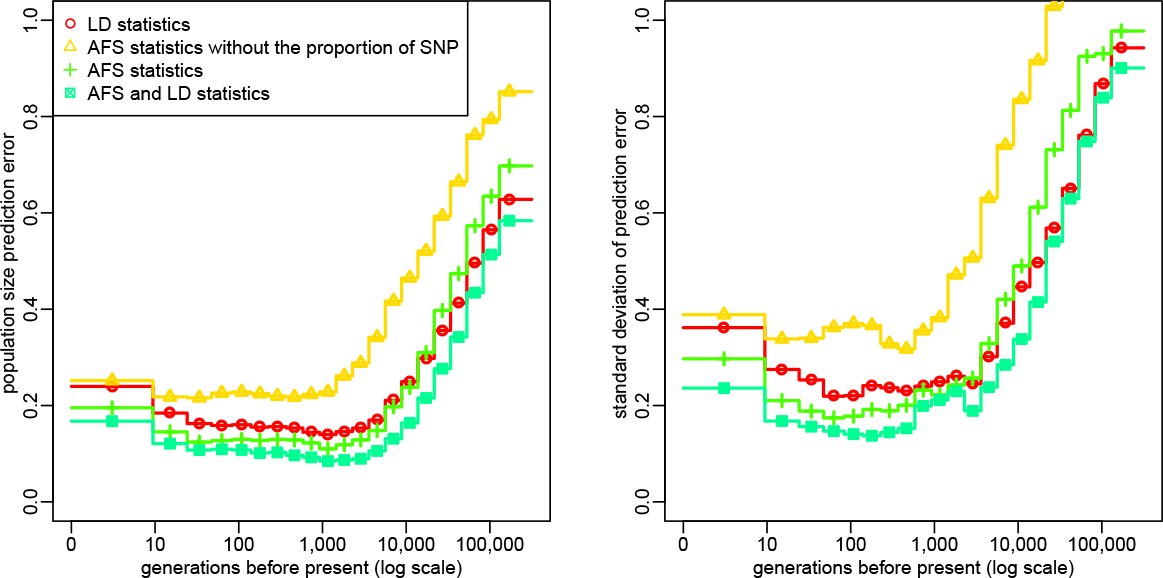
Accuracy of ABC estimation and relative importance of the summary statistics: Prediction error for the estimated population size in each time window (left) and standard deviation of this error (right), evaluated from 2,000 random population size histories. Summary statistics considered in the ABC analysis included different combinations of (i) the AFS (possibly without the overall proportion of SNPs) and (ii) the average zygotic LD for several distance bins. These statistics were computed from *n* = 25 diploid individuals, using all SNPs for AFS statistics and only those with a MAF above 20% for LD statistics. The posterior distribution of each parameter was obtained by neural network regression [32], with a tolerance rate of 0.005. Population size point estimates correspond to the median of the posterior distribution. The prediction errors were scaled in order that point estimates obtained from the prior distribution would result in a prediction error of 1.

### Influence of the amount of data on ABC estimation

Another important question was to assess the amount of data that needs to be simulated and observed in order to achieve optimal accuracy. We first studied the influence of the number of simulated samples and found that increasing this number above 450,000 would not improve the estimation. Indeed, equally low PE and equally small (and accurate) confidence intervals could be obtained using 200,000 simulated samples (S5 Fig).

We then considered the influence of the genome length of observed and simulated samples (S6 Fig). As expected, PEs and the width of credible intervals decreased when the genome length increased. However, only small differences were observed between the performances obtained with 50 and 100 2Mb-long segments, and generating simulated data sets with much more than 100 2Mb-long segments (the default setup considered here) would become very challenging from a computational point of view (see the Methods for more details). For the analysis of observed data sets with a genome length above 200Mb, we thus considered the alternative strategy consisting in comparing observed statistics computed from the full genome (which is computationally very easy) with simulated statistics computed from a subset of the genome. We may think about these simulated summary statistics as an approximation of the genome-wide simulated statistics. To evaluate this strategy, we assumed that the genome length was 100 2Mb-long segments in the observed sample and 10 2Mb-long segments in the simulated samples (S7 Fig). Credible intervals were only slightly improved compared to using a genome length of 10 2Mb-long segments in both simulated and observed datasets, but PEs and their variance between scenarios were reduced, especially for the most recent and the oldest time windows, reaching values almost as low as those obtained when using 100 2Mb-long segments in both simulated and observed datasets. This strategy was thus applied in the further sections of the manuscript, where simulated statistics used for ABC estimation were computed from genomes made of 100 2Mb-long segments, independently of the genome length in the observed data.

We also studied the influence of sample size on population size estimations (S8 Fig). Comparing several sample sizes from n = 10 to n = 50, we observed that using large samples resulted in a more accurate estimation of population sizes in the first 100 generations BP. For instance, in the most recent time window, PE was equal to 0.153 for n = 50 versus 0.212 for n = 10, and the 90% credible interval was narrower (ratio between upper and lower bound of 36 versus 74). These improvements resulted from the fact that low frequency alleles, which are better captured from large samples, are very informative about recent population history. In contrast, population sizes at times more ancient than 10,000 generations BP were more accurately estimated from small samples, although the magnitude of this effect was lower than for recent population sizes (PE of 0.66 for n = 50 versus 0.63 for n = 10 in the most ancient window). This is likely due to statistical overfitting: increasing the sample size leads to increasing the number of AFS statistics, so if these additional statistics are not sufficiently informative they may introduce some noise and reduce the prediction ability of the model.

Finally, we found that computing AFS statistics only from SNPs exceeding a given minor allele frequency (MAF) threshold (from 5 to 20%) resulted in larger PEs and confidence intervals, except for the most ancient population sizes (S9 Fig). Again, this comes from the fact that low frequency alleles are very informative about recent population history. However, as we discuss later, introducing a MAF threshold might be necessary for the analysis of real data sets, so it is interesting to note that even with a MAF threshold of 20% the PE was not much larger than with all SNPs (0.24 versus 0.17 in the worst case).

### Estimation of specific demographic scenarios using ABC

To illustrate the performance of our ABC approach, we then considered six specific demographic scenarios: a constant population size of 500, a constant population size of 50,000, a population size declining from 40,000 to 300 individuals between 3,600 and 100 generations BP, a population size increasing from 2,500 to 60,000 individuals between 1,500 and 250 generations BP, a population size experiencing one expansion from 6,000 to 60.0 individuals followed by a bottleneck of the same magnitude, between 34.0 and 900 generations BP, and a “zigzag” scenario similar to the previous one but including one additional bottleneck between 520 and 50 generations BP (see Fig. 3 for more details). The decline scenario was chosen to mimic the estimated population size history in Holstein cattle [12], the expansion scenario was chosen to mimic the estimated population size history in CEU humans [10], and the “zigzag” scenario has been proposed in [10] as a typical example of very complex history. For each scenario, we simulated 20 PODs of 25 diploid genomes, each genome consisting in 500 independent 2Mb-long segments.

**Figure 3.**
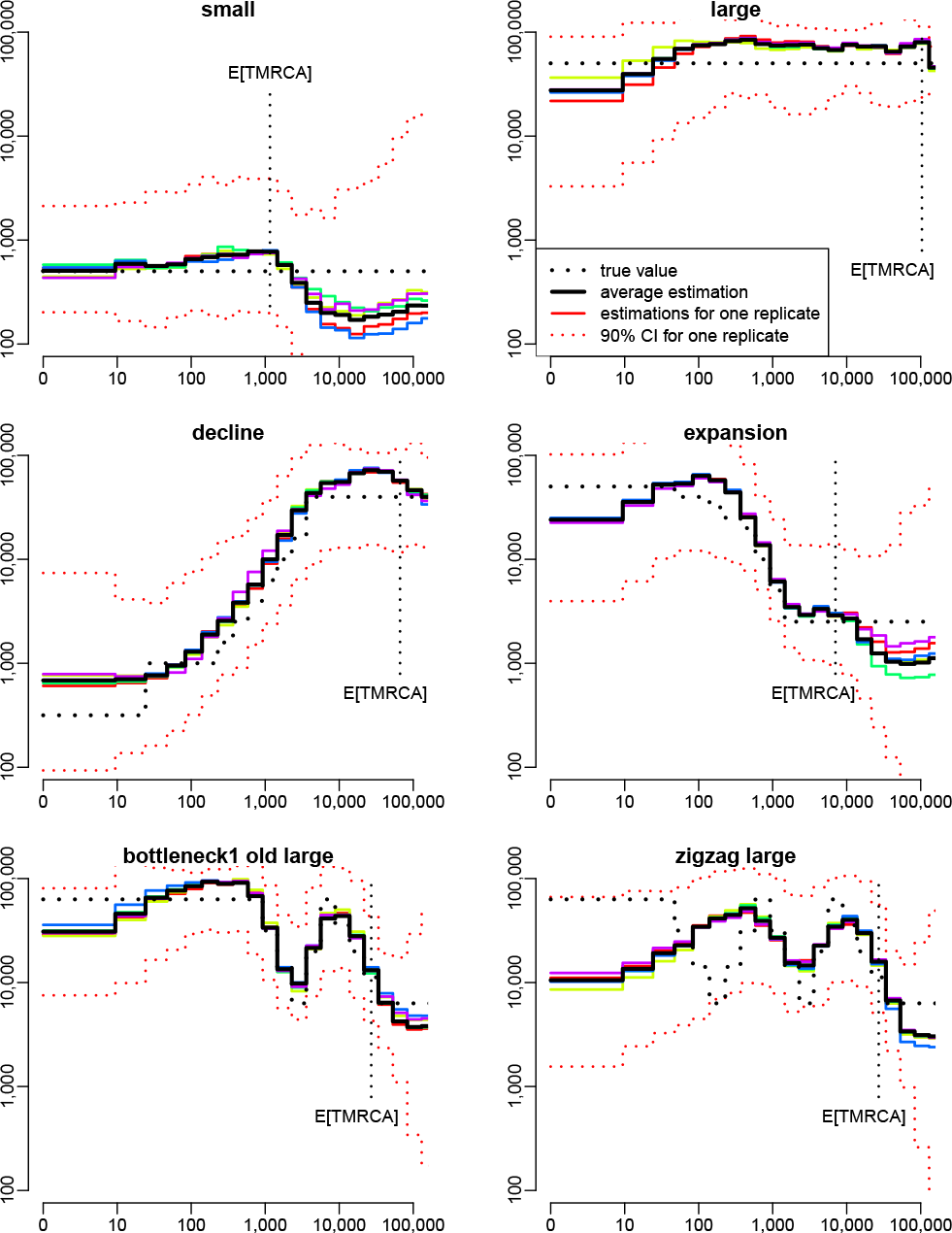
**Estimation of population size history using ABC in six different simulated scenarios:** a small constant population size (*N* = 50,000, top left), a large constant population size (*N* = 50,000, top right), a decline scenario mimicking the population size history in Holstein cattle (middle left), an expansion scenario mimicking the population size history in CEU human (middle right), a scenario with one expansion followed by one bottleneck (bottom left) and a zigzag scenario similar to that used in [10] (bottom right), with one expansion followed by two bottlenecks. For each scenario, the true population size history is shown by the dotted black line, the average estimated history over 20 PODs is shown by the solid black line, the estimated histories for five random PODs are shown by solid colored lines, and the 90% credible interval for one of these PODs is shown by the dotted red lines. The expected time to the most recent common ancestor (TMRCA) of the sample, *E[TMRCA]*, is indicated by the vertical dotted black line. Summary statistics considered in the ABC analysis were (i) the AFS and (ii) the average zygotic LD for several distance bins. These statistics were computed from *n* = 25 diploid individuals, using all SNPs for AFS statistics and SNPs with a MAF above 20% for LD statistics. The posterior distribution of each parameter was obtained by neural network regression [32], with a tolerance rate of 0.005. Population size point estimates were obtained from the median of the posterior distribution.

We observed that all PODs from a same scenario provided very similar ABC estimations (Fig. 3). This suggests that increasing the observed genome length would not improve the obtained estimations, at least with the levels of mutation (1e-8 per bp) and recombination (5e-9 per bp) and the population sizes considered here. Besides, as expected from our previous simulation results, population size history could be reconstructed for all scenarios from a few generations BP back to at least the expected TMRCA of the sample, with the only two exceptions described below.

First, population size estimations in the most recent time window (less than 10 generations BP) often showed a slight bias towards intermediate values, as can be seen in the large constant size scenario, the decline scenario and the expansion scenario. This partly comes from the fact that we estimated population size by the median of the posterior distribution, which tends to shrink it away from our prior boundaries. When estimating population sizes from the mode of the posterior distribution, we were able to better reconstruct the very recent population size in these three scenarios (S10 Fig). Nevertheless, using the mode also brought other issues: it led to less smooth population size histories (S10 Fig) and, on average, to larger PEs than using the median (Fig. 1). Second, the zigzag scenario was incompletely reconstructed: the initial increase of population size and the subsequent first bottleneck could be recovered, but the second bottleneck was replaced by a slow decline.

In order to explore why ABC failed to fully reconstruct this zigzag history, we considered five variants of this scenario (S11 Fig). For a zigzag scenario with smaller population sizes than the original one (ten times lower in all time windows), we observed that ABC could recover the full sequence of expansions and contractions (S11 Fig, top right). This was also the case when only one of the two bottlenecks of this “zigzag small” history was simulated (S11 Fig, bottom). In contrast, when only the most recent bottleneck of the “zigzag large” scenario was simulated, ABC could still not reconstruct it (S11 Fig, middle left). Actually, the decline wrongly estimated by ABC in this case led to very similar summary statistics as the true bottleneck (S12 Fig), and the population size trajectory corresponding to the true bottleneck was included in the 90% credible interval inferred by ABC (S11 Fig, middle left). We also observed that PODs simulated under the wrong decline history would lead to very similar ABC estimations that those simulated under the true bottleneck history (S11 Fig, middle, left vs right). These results suggest that the accuracy of our ABC approach is not strongly affected by the complexity (i.e. the number of expansions and declines) of the true history, but that some specific demographic events, in particular those implying recent population size changes in large populations, can be difficult to identify using this approach. This conclusion was supported by the study of four additional complex scenarios, implying similar expansions and declines as in S11 Fig but in a different order, i.e. the first event was a bottleneck and it was followed by a population decline (S13 Fig). Except the recent part of the “bottleneck2 recent large” scenario (S13 Fig, top left), all aspects of these histories occurring more recently than the expeted TMRCA were accurately reconstructed by ABC.

Because one of our objectives was to estimate the population size history in taurine cattle, we studied more precisely the continuous decline scenario that is expected in this species [12], and evaluated if variations from this scenario could be detected by ABC (S14 Fig). We found that a decline of the same magnitude (from 40,000 to 300), but occurring suddenly either 200 generations BP (top right) or 1,000 generations BP (middle left), would lead to a clearly distinct ABC estimation, although ABC had a tendancy to smooth population size changes. We also considered two scenarios where population size increased again after the sudden decline occuring 1,000 generations BP, either quickly to a relatively high value (5,000, middle right) or more recently to a lower value (1,000, bottom left). In the two scenarios, both the bottleneck phase and the recovery phase could be inferred by ABC. Finally, we studied an alternative scenario where the initial continuous decline was followed by a sudden decline to 100 between 230 and 140 generations BP and by a later recovery to 1,000 (bottom right). Assuming generation time in cattle is about 5 years, the time frame of this bottleneck (between 1,150 and 700 years BP) would correspond to the Middle Age period, where cattle population sizes may have decreased drastically because of wars, famines and cattle plagues [34]. Again, we found that ABC should be able to distinguish this scenario from a simple continuous decline.

### Comparison with MSMC

For each scenario of Fig. 3, we also analyzed five simulated samples with MSMC [10], using two, four or eight of the haplotypes from each sample. When applied to two haplotypes, MSMC is an improved version of PSMC [9], a software that has been used to estimate population size history in many different species within the last few years [35-38]. In our simulations, MSMC based on two haplotypes provided a very accurate estimation of the population size history within a time window starting between a few hundreds and a few thousands generations BP, depending on the scenario, and finishing after the expected TMRCA of 50 haplotypes (Fig. 4). Within this time window, estimations obtained by MSMC from the five replicates were all very close to the true history, even more than ABC estimations. Outside this window however, population size histories estimated by MSMC often had a totally different trend than the true history, (see for instance the small constant size or the decline scenario), with large differences observed between samples (see for instance the expansion scenario). Similar results were obtained when using MSMC with four (S15 Fig) or eight (S16 Fig) haplotypes, except that the time window where accurate population sizes could be obtained was shifted towards recent past, as already shown in [10]. This comes from the fact that MSMC inference is based on the time to the most recent coalescence event, which decreases when the sample size increases. Thus, reconstructing the entire population size history is generally not possible from a single MSMC analysis. This would require concatenating different parts of the history estimated independently using different sample sizes, which might be quite challenging in a real data analysis, because the bounds to consider for such a concatenation are unknown. Besides, for four out of the six scenarios (the small constant size, the expansion, the bottleneck and the zigzag), population sizes at times more recent than approximately 100 generations BP could not be estimated by any MSMC analysis. Indeed, the analysis with eight haplotypes, which is expected to be the most accurate for reconstructing recent demography, provided unstable results for these scenarios. Several other cases where MSMC failed to reconstruct properly the recent history were observed among the additional scenarios tested with ABC, as for instance in the “bottleneck cattle middle age” scenario (S17 Fig and S18 Fig) for which the recent bottleneck was not detected.

**Figure 4.**
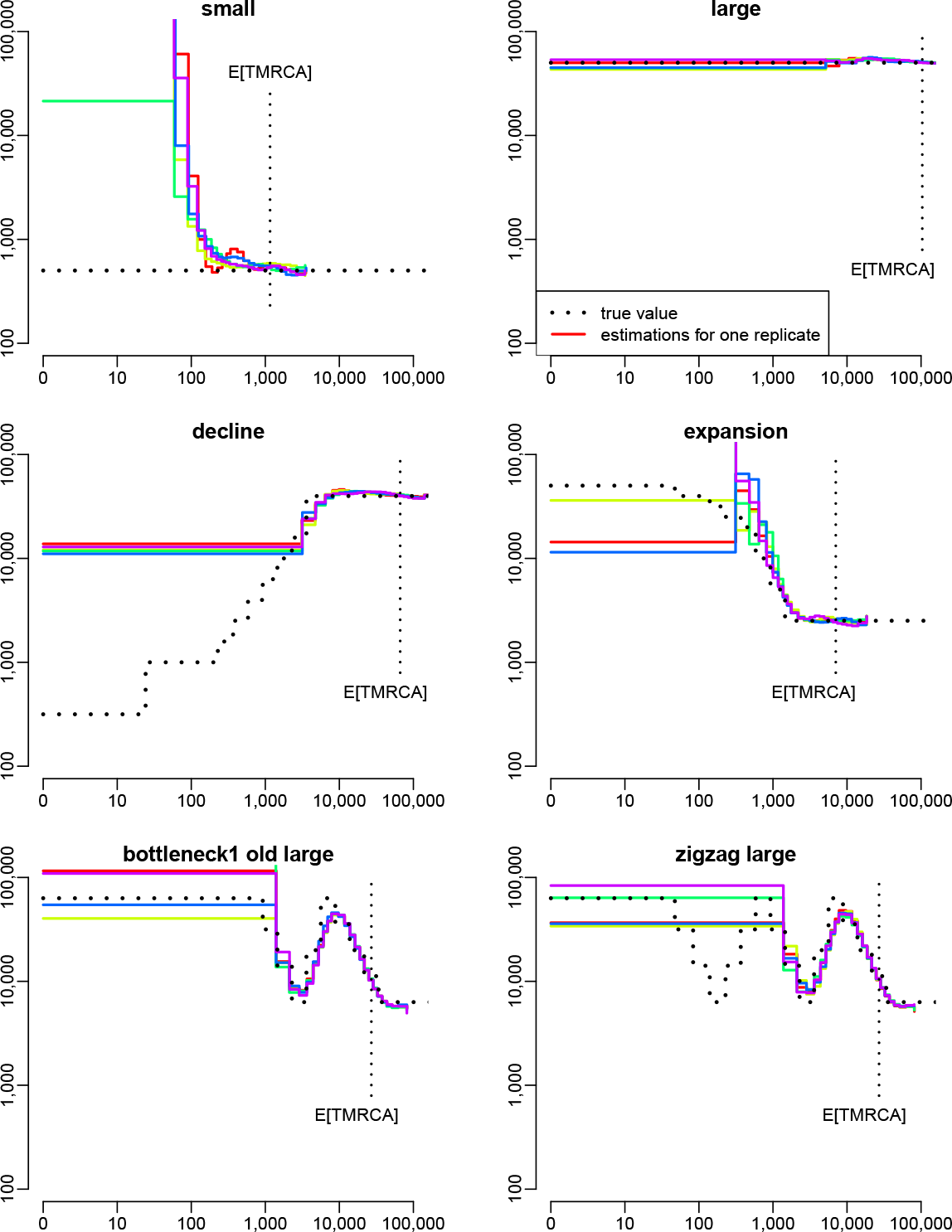
Estimation of population size history using MSMC with two haplotypes in five different simulated scenarios: For each scenario, the five PODs considered for MSMC estimation were the same as in Fig. 3. The expected TMRCA shown here is also the same as in Fig. 3, it corresponds to samples of 50 haploid sequences.

Finally, it is important to note that the simulated data that we used in these MSMC analyzes were assumed to be perfectly phased. However, real data consist generally in statistically inferred haplotypes, which can typically include from 1 to 10 switch errors per Mb and individual, even when using recent phasing algorithms and large sample sizes [39]. In our simulations, analyzing phased data with such switch error rates often biased MSMC estimations, especially for the most recent part of the demographic history (S19 Fig). To avoid this issue, MSMC can in principle be run from unphased data, but we found that this would also affect the estimation accuracy (S19 Fig, right column).

### Application to NGS samples in cattle

We applied our ABC approach to estimate the population size history in four cattle breeds, using large samples of diploid genomes recently published by the 1,000 bull genomes project [40]. An important issue when analyzing NGS data is the potential influence of sequencing and phasing errors on the estimations. To investigate this question, we first evaluated how these errors affect the summary statistics considered in our ABC approach. We considered a set of 12 Holstein animals for which the haplotypes inferred from NGS data within the 1,000 bull genomes project could be compared with those inferred from 800K SNP chip data obtained independently from another project. Assuming that 800K data are free of genotyping errors, we computed the summary statistics from these data and checked whether similar values could be obtained from NGS data at the same positions (S20 Fig). We found that the average gametic LD (i.e. the LD computed from haplotype data) was significantly smaller with NGS data than with 800K data at long physical distances, but not at short ones. This likely comes from an increased level of phasing errors in NGS data as compared to 800K data. Indeed, such errors tend to artificially break the correlation between SNPs within each individual, which reduces LD. Besides, as they are relatively rare, we expect their influence to be significant only when comparing SNPs at large physical distance.

In contrast, the average zygotic LD (i.e. the LD computed directly from genotype data) was identical for the NGS and the 800K data. We also observed a perfect match between the polymorphic site AFS obtained from the NGS data subsampled at 800K positions, and from 800K data. Finally, the overall proportion of SNPs was similar in the two types of data. More precisely, based on the 800K positions and the sample of 12 individuals, we found approximately 0.5% of false positive SNPs, i.e. positions that were found polymorphic when using NGS data but not when using 800K data (S21 Fig, left), and approximately 5% of false negative SNPs, i.e. positions that were found polymorphic when using 800K data but not when using NGS data (S21 Fig, right). Besides, the proportion of false negative SNPs did not depend on the true allele frequency (i.e. the allele frequency in the 800K data), so it should not distort the AFS. Overall, these results suggest that our summary statistics, when computed from genome wide unphased NGS data, should not be affected by sequencing and phasing errors. However, the above comparison does not really allow to evaluate the influence of false positive SNPs when analyzing genome wide NGS data, because the 800,000 positions of the SNP chip are strongly enriched in true SNPs compared to the three billions of positions of the entire genome.

To overcome this limitation, we studied directly the influence of sequencing and phasing errors on ABC estimations, by analyzing one sample of 25 Holstein genomes with slightly different combinations of summary statistics (Fig. 5). When LD was computed from haplotypic data, the estimated recent population size was above 20,000 individuals, which seems quite unrealistic given that the estimated current effective size of this breed is generally of an order of 100 [17-19, 41]. This discrepancy likely resulted from the average LD at large physical distances, which was artificially reduced by phasing errors, as discussed above. Computing LD from genotypic data, we obtained more realistic results, with a recent population size of 7,000. However, there was a great difference between the estimation obtained when computing AFS statistics from all SNPs, and that obtained when computing these statistics only from SNPs with a MAF above 10% (Fig. 5). Such a large difference was not expected from simulations, neither on average over multiple random histories (S9 Fig, middle) nor in the particular cases of a constant or declining population (Fig. 3 vs S22 Fig). Thus, it must result from the influence of false positive SNPs, which are much more likely to produce low frequency alleles (S21 Fig, left). In contrast, there was little difference between the estimations obtained when computing AFS statistics with a MAF threshold of 10 or 20%, which strongly suggests that these strategies are both robust against sequencing errors, at least for this particular dataset. To be conservative, we used a MAF threshold of 20% for the final analysis of the four breeds.

**Figure 5.**
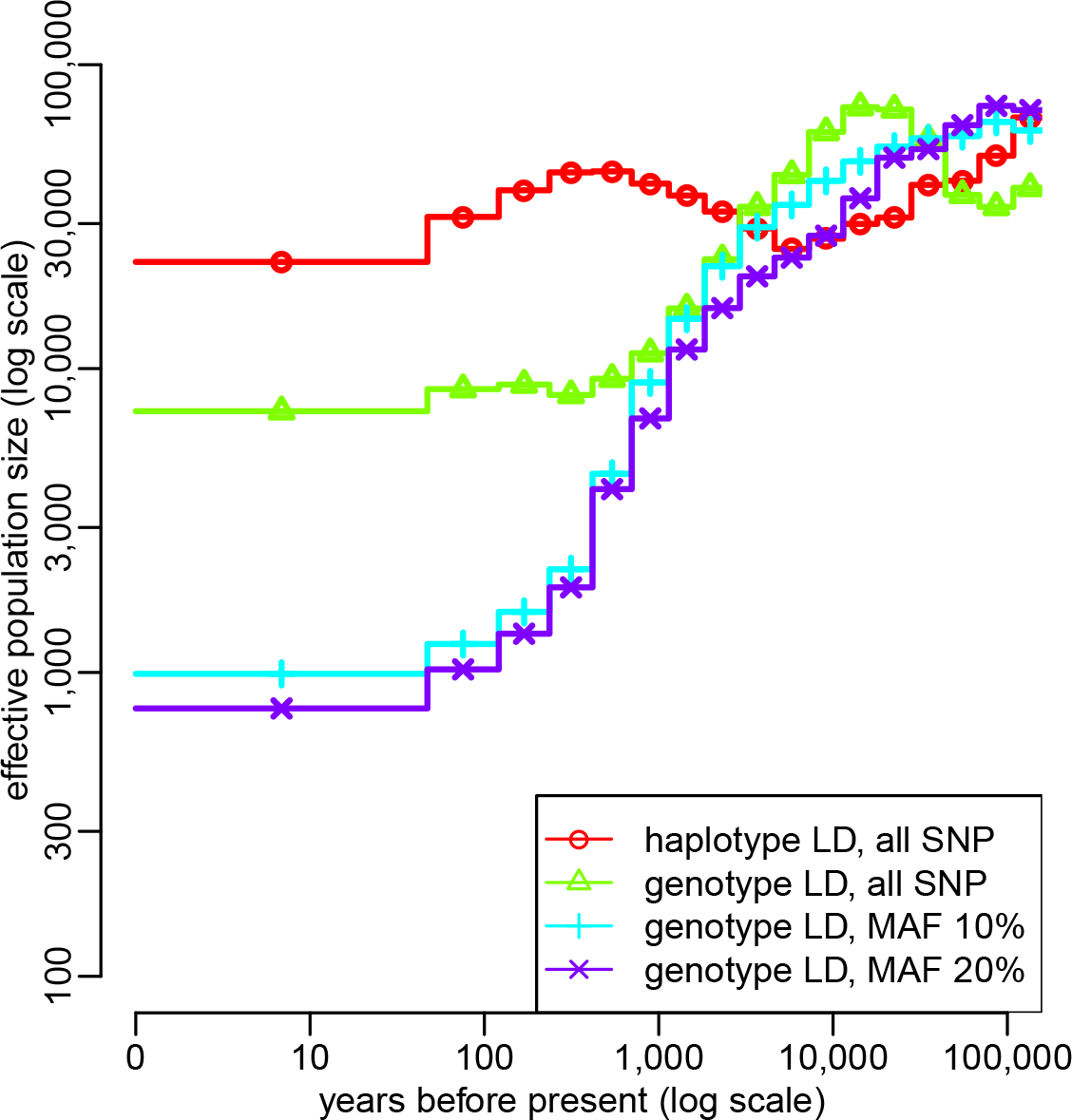
Influence of phasing and sequencing errors on ABC estimation: Estimation of population size history in the Holstein cattle breed using ABC, based on whole genome NGS data from *n* = 25 animals. Summary statistics considered in the ABC analysis were (i) the AFS and (ii) the average LD for several distance bins. LD statistics were computed either from haplotypes or from genotypes, using SNPs with a MAF above 20%. AFS statistics were computed using either all SNPs or SNPs with a MAF above 10 or 20%. The posterior distribution of each parameter was obtained by neural network regression [32], with a tolerance rate of 0.005. Population size point estimates were obtained from the median of the posterior distribution. Generation time was assumed to be five years.

This analysis outlined several interesting features of cattle demographic history (Fig. 6). Before 10,000 years BP, the population sizes estimated in the four breeds were very similar, in agreement with the fact that all four breeds descend from a same ancestral population, i.e. the initial *Bos taurus* population which resulted from the domestication of the wild aurochs, *Bos primigenius*, approximately 10,000 years BP [42]. This common estimated history is characterized by a population decline starting approximately 50,000 years BP. In particular, a sharper decrease was observed from approximately 20,000 years BP, which could correspond to the intensification of anthropogenic effects like hunting or later herding [42]. Shortly after domestication, the inferred population size histories could be divided into two groups, Holstein and Fleckvieh on one hand, Angus and Jersey on the other hand. This is consistent with the origin of these breeds: Holstein and Fleckvieh ancestors were brought into Europe through the *Danubian route* approximately between 7,500 and 6,000 years BP, while Angus and Jersey have more diverse origins and partly descend from animals that were brought into Europe through the *Mediterranean route* approximately between 9,000 and 7,300 years BP [43,44]. Population size histories in the four breeds finally diverged during the last 500 years, which is consistent with the progressive divergence of these breeds induced by geographic isolation and, from the 18th century, by the creation of modern breeds [45]. This lead to recent effective population sizes of 290 in Angus, 390 in Jersey, 790 in Holstein and 2,220 in Fleckvieh.

**Figure 6.**
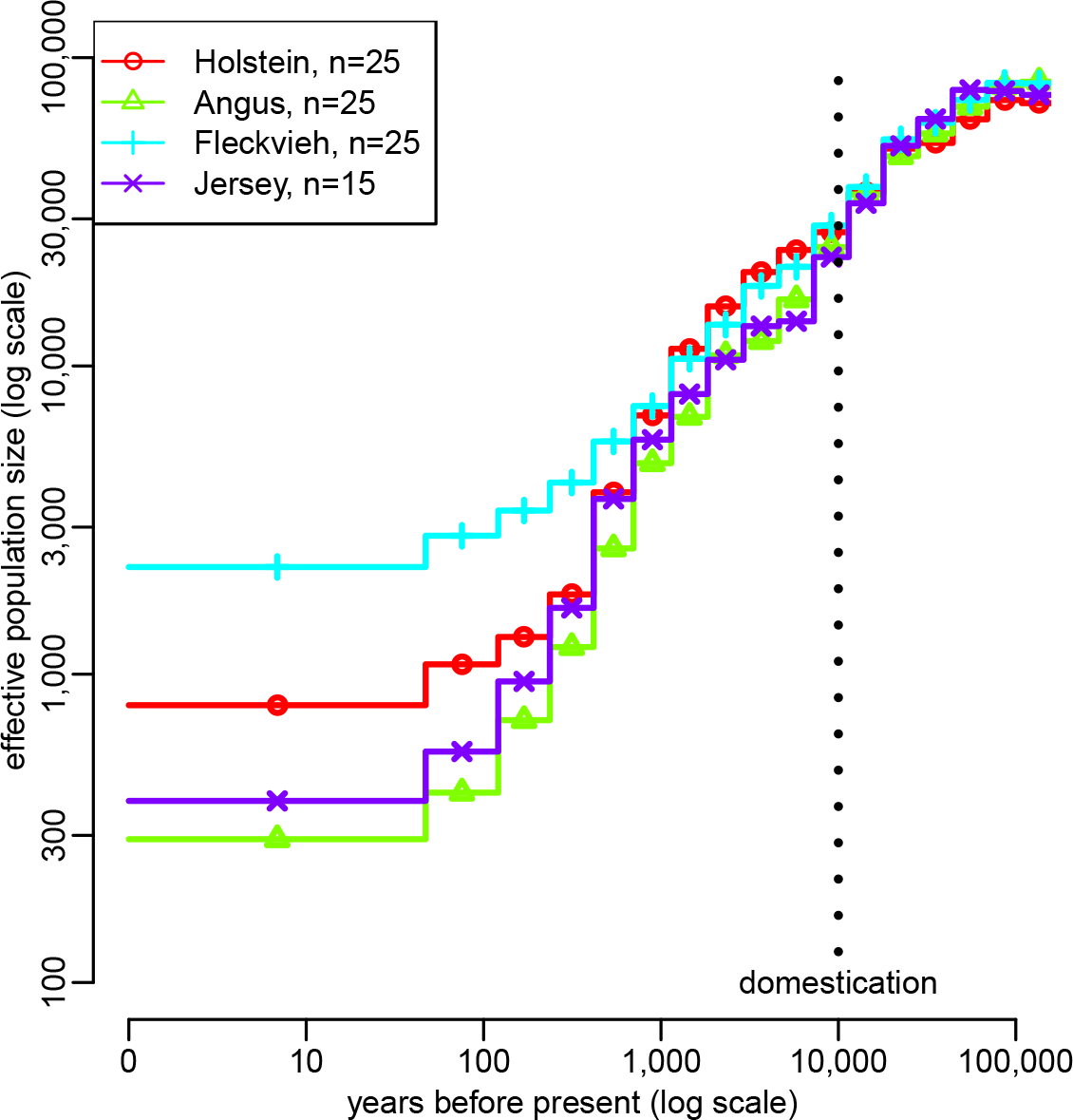
Estimation of population size history in four cattle breeds using ABC: Angus *(n* = 25 animals), Fleckvieh (*n* = 25), Holstein (*n* = 25) and Jersey (n = 15). Estimations were obtained independently in each breed, based on whole genome NGS data from sampled animals. Summary statistics considered in the ABC analysis were (i) the AFS and (ii) the average zygotic LD for several distance bins. These statistics were computed using SNPs with a MAF above 20%. Other parameter settings are the same as in Fig. 5.

The 90% credible intervals associated to these estimated population size histories are shown in S23 Fig. We performed posterior predictive checks by sampling population size histories from the posterior distributions and simulating new genomic samples from these histories [46]. The summary statistics obtained from these samples were similar to those observed in the real data (S24 Fig). We also checked that the best simulated histories provided summary statistics that were indeed similar to the observed summary statistics (S25 Fig). Finally, we note that point estimations of the average per site per generation recombination rate were quite similar between breeds: it was equal to 3.66e-9 in Holstein, 3.89e-9 in Fleckvieh, 4.58e-9 in Jersey and 5.00e-9 in Angus.

## Discussion

### Methodological contribution

Applying our ABC approach to genomic samples simulated under a large number of random population size histories, we showed that it provides, on average, accurate estimations of population sizes from the first few generations BP back to the expected TMRCA of the sample. Because the estimation accuracy depends on the true population size history, we also analyzed genomic samples simulated under 20 specific demographic scenarios with various levels of complexity: a constant population size (2 scenarios), a monotonic decrease (3 scenarios) or expansion (1 scenario), a single bottleneck (3 scenarios), a single bottleneck plus an additional expansion or decrease (9 scenarios) or two bottlenecks plus an additional expansion (2 scenarios). For most of these scenarios, PopSizeABC could reconstruct the population size history from present time back to the expected TMRCA of the sample. Within this time limits, the only situations where the ABC point estimates were very different from the true history were those implying a decline or expansion occurring in a large population (more than 5,000 individuals) within the last few hundreds generations. Indeed, when large population sizes are combined with frequent population size changes (in our model, recent time windows are also the shortest ones), each time window represents a very small part of the coalescent history, which explains why these scenarios are particularly difficult to reconstruct. However, in these situations, the true history was still included within the 90% credible interval, and the increased width of this interval compared to other time windows suggested that the point estimate was less reliable. Similarly, in all scenarios, the width of the credible interval increased rapidly for times that were more ancient than the expected TMRCA, which corresponds thus to the upper bound of the time period where ABC estimation could be trusted.

Interestingly, we observed that PopSizeABC behaved quite differently from MSMC [10], a recent full-likelihood SMC-based method allowing to analyze multiple diploid genomes. On one hand, for the 20 scenarios considered here, MSMC estimated more accurately than PopSizeABC the population sizes at several time points. This was expected because ABC inference implies a much larger degree of approximation than MSMC inference. On the other hand, the total time period for which each demographic history could be correctly reconstructed with a single MSMC analysis was much smaller than with ABC. Besides, in most scenarios, recent population sizes (in the first 100 generations BP or even more) could not be inferred by any MSMC analysis, while they could be inferred by ABC. In our study of cattle demography, reconstructing the population size history for this recent period allowed to highlight the specificity of each breed. In many other situations, and especially in a conservation perspective, estimating recent demography is actually crucial.

The better performance of ABC to reconstruct recent population size history is partly explained by the possibility of using larger samples. We generally considered samples of 25 diploid genomes, which resulted in more accurate estimations of population sizes in the last 30 generations than using only 10 diploid genomes (S8 Fig). Indeed, large samples contain rare alleles. Since these alleles result from mutations that occurred in the most recent part of the coalescent tree, their relative proportion in the AFS is informative about the recent variations of population size. Interestingly, gaining accuracy for recent time periods by increasing the sample size had no strong negative impact on the reconstruction of the older demographic history (except for times older than the TMRCA), contrary to what was observed with MSMC. The use of LD statistics must also contribute to the reconstruction of recent demography because, in our simulations, predictions of population sizes at times more recent than 100 generations BP were still acurate when rare alleles were removed (S9 Fig). As discussed below, the average LD at long physical distances is expected to reflect the recent population size [26].

Following previous studies [26, 30, 47], we used in our ABC approach the average LD over different bins of physical distance in order to get information about population sizes at different times in the past. In a finite population, LD results from a balance between drift and recombination. This implies that LD between markers at long recombination distance mostly reflects recent population sizes, while LD at short recombination distance also reflects ancient population sizes [48]. To illustrate this, we computed our LD statistics for several simulation scenarios consisting in a sudden expansion with fixed magnitude but occurring at different times in the past (S26 Fig, left). As expected, we observed that LD statistics at long distance were similar to those of a large population, thus reflecting the recent population size, while LD statistics at small distance were similar to those of a small population, thus reflecting the ancient population size. Besides, the more recent the expansion, the larger the distance required to observe a LD level reflecting the large (recent) population size. Similarly, for decline scenarios, markers at long (resp. short) distance were most of the time found to reflect the LD level in a small (resp. large) population (S26 Fig, right; see the legend for more details)).

This relation between the recombination distance and the time horizon can even be described more precisely. If population size is assumed to change linearly over time, it can be shown that the expected *r*^2^ between SNPs at recombination distance c is approximately equal to

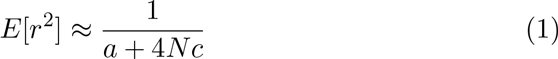

where N is the effective population size at time 1/(2*c*) BP and *a* is a constant depending on the mutation model [26]. The evolution of population size through time can thus be reconstructed by computing the average *r*^2^ for different bins of recombination distance, and then inverting the formula in (Eq. (1) [26,47]. However, several authors pointed out that this approximation is unsatisfactory, especially for non constant demography [49,50], and could lead to wrong estimations of past population sizes [50,51]. Our ABC approach overcomes this issue, because *r*^2^ values estimated from the data are not compared to approximate theoretical predictions, but to simulated *r*^2^ values. Using this approach, we could demonstrate that these statistics contain useful information about the population size history (Fig. 2). We further demonstrated two important properties of LD statistics in the context of population size inference (S4 Fig). First, computing *r*^2^ from genotypes is as informative as computing it from haplotypes, in the sense that it leads to similar PEs. Second, removing rare SNPs (at least those with MAF below 5%) when computing this LD measure reduces PE.

In our simulations, ABC inferences based on AFS statistics alone also provided accurate estimations of population sizes at different times in the past (Fig. 2). Theoretical studies have demonstrated that complex population size histories can be estimated from AFS statistics [52], and these statistics are already the basis of several inferential approaches in population genetics [13,27-29,31]. In particular, two recent studies implemented composite-likelihood approaches to estimate population size through time in a single population [13,31], and obtained convincing results on simulated data. We do not expect that our ABC approach based on AFS statistics alone would improve the point estimations obtained by these approaches, and analyzing very large samples (i.e. hundreds or thousands of individuals) would certainly be much more challenging with ABC due to the simulation step. However, one advantage of ABC is to provide credible intervals, which allow to quantify the degree of confidence associated to a given point estimation.

Moreover, one important conclusion of our work is that combining AFS and LD clearly improves, on average, the estimation of population sizes (Fig. 2). This stems from the fact that these two classes of statistics are not informative for the same demographic scenarios. While prediction errors obtained from AFS or LD statistics were quite similar for scenarios with little population size variations (S27 Fig, top panels), better predictions were obtained from AFS (resp. LD) statistics when the main trend of the population size history was an expansion (resp. a decline) (S27 Fig, bottom panels). These differences were mainly due to the predictions obtained from AFS statistics, which were much better for expansion scenarios than for decline scenarios. Indeed, population declines accelerate the rate of recent coalescence events compared to old ones. Combined with the fact that the time intervals between recent coalescence events are intrinsically shorter than between old ones (because coalescence rates are proportional to the square of the sample size), this tends to produce coalescence trees where only the few oldest branches have a substantial length. In other words, the recent topology of coalescence trees in decline scenarios is very hard to infer based on observed data, making it difficult to estimate population size variations from the AFS. These results are consistent with those from a recent study [53], which showed that, for the inference of single bottleneck events, including some linkage information was more efficient than using the AFS alone. Actually, one interesting conclusion of S27 Fig is that combining LD and AFS statistics always improves the prediction compared to using either one or the other class of statistics alone, whatever the family of scenarios we considered.

This conclusion was also supported by the study of several specific scenarios: some could be accurately reconstructed from AFS statistics alone but not from LD statistics alone (Figure 7, top), and vice versa (Figure 7, middle), but the prediction obtained when combining AFS and LD statistics was always close to the best of the two. In other scenarios, neither AFS or LD statistics alone allowed to correctly estimate the demographic history, and using them jointly was therefore essential (Figure 7, bottom). Finally, in many scenarios, ABC estimation based either on LD statistics alone or AFS statistics alone performed already very well, but the advantage of combining AFS and LD statistics clearly appeared when using a MAF threshold that reduced the information brought by AFS statistics (S28 Fig). Besides these effects on population size estimation, note that combining AFS and LD statistics allowed to estimate the average per site recombination rate, which was not possible using either one or the other class of statistics alone.

**Figure 7.**
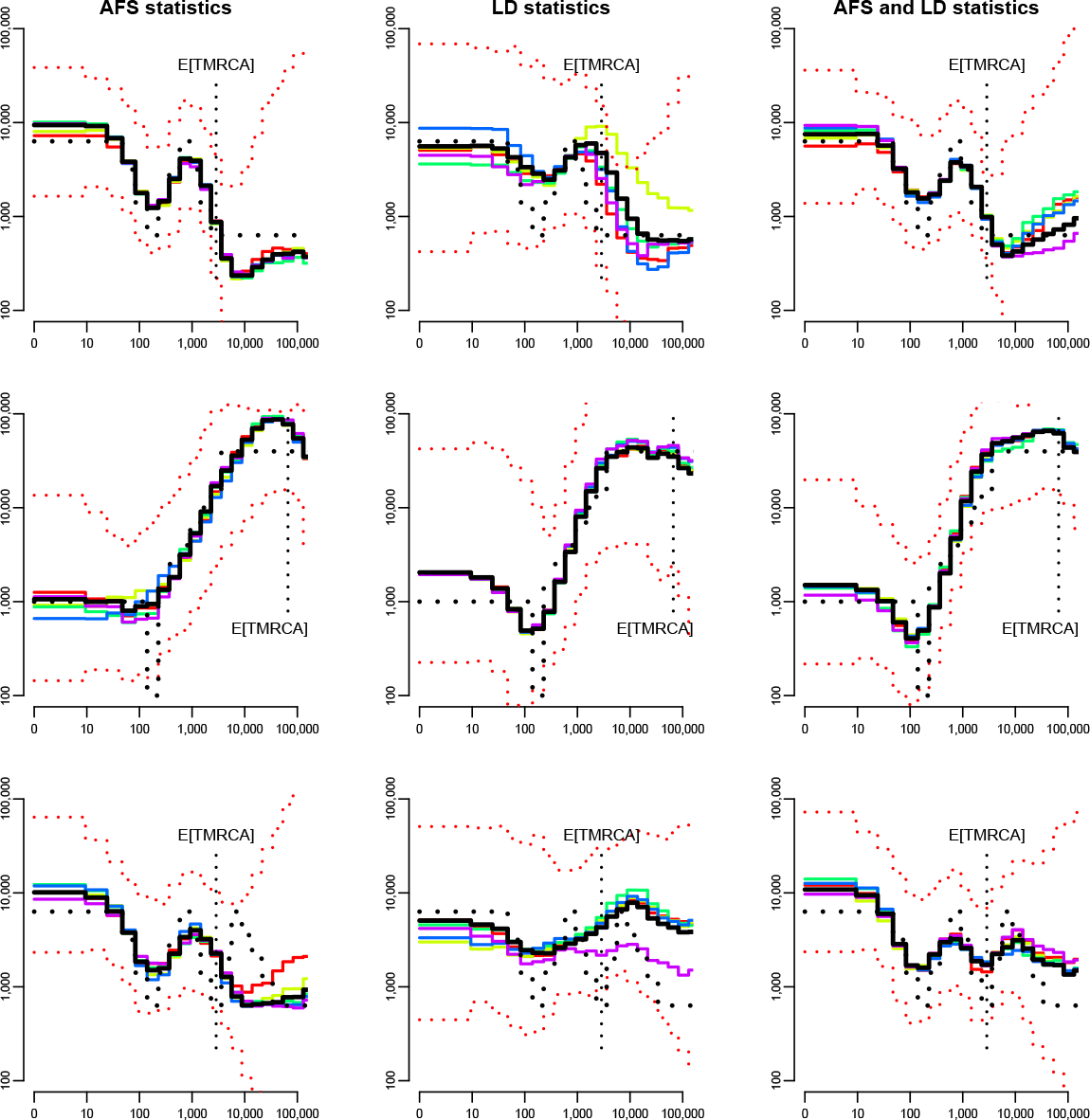
Comparison of summary statistics for the estimation of population size history in three scenarios: “bottleneckl recent small” (top), “bottleneck cattle middle age” (middle) and “zigzag small” (bottom). Summary statistics considered in the ABC analysis were either the AFS statistics alone (left column), the LD statistics alone (middle column), or the AFS and LD statistics together (right column). All other settings are similar to Fig. 3, as well as the legend.

The genome wide distribution of the length of IBS segments shared between two chromosomes could provide another interesting class of summary statistics for ABC, because several recent studies showed that it is very informative about population demography [12,54]. However, we found that applying ABC from a set of statistics related to this distribution, rather than from AFS and LD statistics, resulted in larger PEs of population sizes more recent than 100 generations BP (S29 Fig). This is likely due to the much smaller number of individuals simultaneously considered in IBS statistics. When IBS statistics were used in addition to AFS and LD statistics, no significant improvement was observed compared to the combination of AFS and LD statistics. Besides, the estimation of recent population demography is mainly influenced by the frequency of long IBS segments, which might be difficult to estimate in practice due to sequencing errors [12,54]. Thus, we did not further investigate the inclusion of these statistics in our approach.

Several previous studies implemented ABC approaches based on genomewide data to infer population genetics models [21-25]. However, none of these studies focused on the estimation of population size through time using complex step-wise models, as we did here. In a Bayesian perspective, this specific question had, so far, only been adressed using a small number of independent non-recombining loci [5-8]. Another originality of our study is to use LD summary statistics that can only be computed from relatively long DNA sequences (at least 2Mb) with recombination, while almost all previous genome-wide ABC studies (but see [23]) considered short loci (≤ 20kb long). Even with modern computer facilities, simulating hundreds of thousands of long DNA sequences required some optimization adjustments. One of them was to reduce the space of possible simulated histories to the most realistic ones by setting constraints on the prior distributions of population sizes (see Methods). Another one was to allow simulated and observed samples to differ in two different ways. First, the total genome length was generally smaller in simulated samples than in the observed sample, which resulted in lower prediction errors than reducing the genome length in the observed sample down to the one that could be efficiently achieved in simulated samples (S7 Fig). Second, when analyzing the cattle data, the simulated summary statistics were computed from independent 2Mb-long segments, although the observed ones were computed from contiguous 2Mb-long segments. Indeed, simulating data under the coalescent with recombination becomes extremely difficult for long sequences. This second approximation cannot bias the estimations, because the correlation structure between segments has no impact on the expected value of summary statistics. Similar to the genome length, the correlation structure of the genome only affects the precision (i.e. the estimation variance) of summary statistics. Despite of the additional correlation, computing summary statistics in cattle using the entire genome (≈ 1,250 contiguous 2Mb-long segments) likely resulted in a higher precision, and thus in a more acurate estimation, than using a subset of 100 independent 2Mb-long segments.

Analyzing real data sets with our approach presents several important advantages. First, our approach is designed to be applied to totally unphased data. Indeed, AFS statistics are deduced from the allele frequencies at all SNPs, which can be computed directly from genotypes. LD statistics are also computed from genotypes, although the common practice in population genetics is to compute them from haplotypes. LD statistics computed from genotypes are not identical to LD statistics computed from haplotypes, but they lead to similar estimations of population sizes. As observed in the analysis of the cattle data (Fig. 5), phasing errors can have dramatic effects on the estimated histories, and they would certainly affect the inference for all populations where the experimental design prevents from phasing the data with high accuracy. Moreover, the SNP data handled by our method can be unpolarized, i.e. it is not necessary to know which of the alleles at a given SNP is ancestral. Using polarized data would probably improve the estimations, as this would allow computing the unfolded rather than folded AFS. However, inferring ancestral alleles is not always possible and is prone to errors, so we chose to focus on statistics computable for all datasets. Finally, based on the analysis of NGS data in cattle, we showed that our approach can easily be made robust to sequencing errors by computing summary statistics only from SNPs with common alleles (MAF ≥ 10 or 20%, Fig. 5). This modification is expected to increase the population size prediction errors and the width of credible intervals if the dataset contains no sequencing errors (S9 Fig), but this seems by far preferable to the large biases caused by sequencing errors, as illustrated by our study and several previous ones [9,12].

One consequence of sequencing errors is to create wrong SNP calls in the data, at genomic positions where the observed sample is actually not polymorphic. Because these wrong SNPs are generally associated to low frequency alleles, focusing on SNPs with common alleles reduces the proportion of wrong SNPs in the data, and consequently their influence on summary statistics. In our application to cattle NGS data, this strategy was efficient because wrong SNP calls were the only detectable effect of sequencing errors on the data. In particular, genotyping errors at true SNP calls had no impact on the summary statistics, as shown by the perfect match between summary statistics computed from NGS data or genotyping data at the 800K chip positions (S20 Fig). Indeed, NGS genotypes had been corrected by imputation, taking advantage of the large sample size and / or sequencing depth within each breed [40]. As this might not be the case in all data sets, other strategies could be applied to correct for sequencing errors, while keeping the main idea of an ABC approach based on AFS and LD statistics. For instance, one could simulate NGS data with the same coverage and error rates as the observed data, rather than perfect genotype data, and compute observed and summary statistics directly from raw NGS data, using dedicated algorithms that account for the uncertainty of genotype calls. Such algorithms are available both for AFS [55] and LD statistics [56], which is another advantage of using these standard summary statistics. However, this strategy would be much more computationally demanding than the one we used here.

### Contribution to the demographic history of cattle

Until recently, effective population size estimations in cattle, and more generally in all livestock species, were mostly based on two approaches. The first approach focuses on the few most recent generations and estimates population size from the increase of inbreeding or coancestry along generations, based on pedigree or molecular information [17-19]. Using this approach, population size estimations from around 50 animals in Holstein to around 150 animals in Simental (closely related to Fleckvieh) were obtained [19]. These estimated populations sizes are qualitatively consistent with ours, as we estimated that the recent population size in Fleckvieh was about three fold larger than in Holstein, but the actual values obtained with these approaches were substantially lower than our estimates (790 in Holstein and 2,220 in Fleckvieh). This may partly be due to the small bias observed with our approach in the simulated decline scenario, using either the median (Fig. 3) or the mode (S10 Fig) of the posterior distribution as point estimation. But it is also important to mention that the animals sequenced in the 1,000 bull genomes project were chosen among key ancestors of the breed, so the most recent population size estimated in our study might reflect the population size a few generations ago rather than the current one. This could partly explain the discrepancy between the estimates, because artificial selection has been particularly intensive within the few last generations, leading to a further decline of effective population size.

The second approach is based on the average *r*^2^ over different bins of genetical distance [26,47], which has been already mentioned earlier in the discussion. It aims at estimating population size on a much larger time scale and has been extensively applied in cattle [41,57,58] and other livestock species [59]. Indeed, a very large number of animals have been genotyped using SNP chips in these species, sometimes for other purposes, such as QTL detection, and used for LD estimation. In addition to the methodological issues related to this approach, the use of SNP chip data for population size estimation presents its own limitations. The ascertainment bias associated to SNP chip data does not only influence AFS statistics but also LD statistics, which in turn affects population size estimation. This is outlined by the fact that our ABC approach based only on LD summary statistics infers different population size histories when these statistics are computed from all the SNPs found by NGS, or only from those that overlap with the 800K chip (S30 Fig). Regrettably, this influence of ascertainment bias on population size estimations obtained from LD statistics is generally not accounted for by the studies using LD. Besides, considering LD alone leads to a different prediction than considering LD and AFS together (S30 Fig), and our simulation results suggest that the former prediction is less reliable. Overall, the use of NGS data, and of dedicated inference approaches taking advantage of these data, should thus considerably improve our understanding of livestock evolutionary history, at least above 600 generations (3,000 years in cattle) BP (S30 Fig).

To our knowledge, the first (and so far the only) estimation of population size history in cattle based on NGS data was obtained by [12]. This result was based on the distribution of IBS segment length in one Australian Holstein bull sequenced at 13X coverage. The overall histories found in this study and in ours are quite consistent, as they both exhibit a strong decline of population size from about 20,000 years BP to the very recent past, but our estimations of population size are generally larger. For instance the population size before this decrease was around 20,000 in their study and around 50,000 in ours, and the population size 1,000 years ago was around 2,000 in their study and around 4,000 in ours. Although the most obvious difference between the two approaches is that they use different summary statistics, ABC estimations obtained from IBS statistics rather lead to larger or equal population sizes than those obtained from AFS and LD statistics (S31 Fig). Thus, we think that the difference between our estimation and that in [12] more likely comes from a difference in the recombination rate. This rate is set to 1e-8 per generation and per bp in [12], while our approach would rather provide an estimation around 4e-9. Assuming that our estimation is correct, the overestimation of r by a factor two in [12] could lead to an underestimation of N by the same factor, because one essential parameter determining the IBS segment length distribution is the scaled recombination rate 2*Nr*. Further work will be needed to better understand the difference between the two estimations.

### Perspectives

Our ABC approach, as well as other SMC [9,10] or IBS based methods [12], assumes that the considered population has evolved forever as an isolated population. This is obviously a strong hypothesis: for instance the cattle breeds considered here have actually diverged from a common ancestral population. Several studies have demonstrated that population structure can leave genomic signatures similar to those of population size changes, even if each of the subpopulations is actually of constant size [60-64]. Consequently, population size histories estimated by single population approaches should be interpreted with caution. However, we anticipate that our study will pave the way for future approaches inferring population size histories jointly in multiple populations, while accounting for the history of divergences and migrations in these populations. ABC represents a perfect framework for developing such approaches, because of the flexibility offered by the simulation procedure. It is already widely used in population genetics for estimating parameters in multiple population models including for instance admixture events and some population size changes [65]. Besides, previous studies showed that structured models and population size change models can be distinguished using ABC [61].

In this study, the flexibility offered by ABC allowed us to infer parameters under the true coalescent with mutation and recombination, rather than under the SMC approximation as in [9,10,54]. One could actually go much further and relax also the hypotheses of the Kingman’s (1982) coalescent. For instance in cattle, genealogies in the most recent generations are highly unbalanced, because a few bulls with outstanding genetic values have been used to produce thousands of offsprings through artificial insemination. Such genealogies are not consistent with the Kingman’s coalescent, but specific algorithms combining the Kingman’s coalescent with a few generations of forward-in-time simulations could certainly be implemented and used to perform ABC estimations in this context.

## Methods

### The ABC approach

Assume we observe a dataset 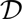, from which we want to estimate the parameters *θ* of a given model. In a Bayesian framework, this involves computing the posterior probability 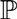(*θ* | 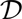) for any possible parameter value. In many situations, and in particular in population genetics, this posterior cannot be derived because of the model complexity and even numerical evaluations are impossible due to the high dimensionality of the observed data space. The idea of ABC [20] is to replace in this context the full dataset 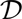 by a vector of summary statistics 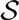 capturing most information contained in the data and to estimate model parameters based on the approximate posterior 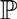(*θ* | 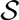). The estimating procedure consists in sampling a very large number of parameter values from a prior distribution, simulating datasets from these parameter values, and accepting the parameter values leading to summary statistics that are sufficiently similar to those of the observed dataset.

Several strategies can then be used to estimate the posterior distribution.

The easiest one, called rejection, is to compute the empirical distribution of the accepted parameter values. To account for the imperfect match between accepted and observed summary statistics, accepted parameter values can also be adjusted by various regression methods, using the associated summary statistics as explanatory variables. The general idea of these methods is to assume a local regression model in the vicinity of *S*, with an equation of the form

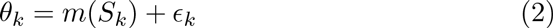

where *θ_k_* is the value of parameter *θ* in the kth simulated sample, *S_k_* is the vector of summary statistics in this sample, *m*() is a regression function varying between approaches, and *ɛ_k_* is a random noise. This model is fitted using all accepted samples. Adjusted parameter values are then obtained by

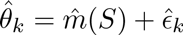

where 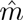 is the estimated regression function and 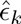is the empirical noise, and the posterior distribution is finally computed as the empirical distribution of these adjusted values. A general review on these aspects can be found in [46].

### Model and priors

Here the observed data 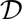 is a set of *n* diploid genomes sampled from a single panmictic population, and the model assumed to have generated these data is the coalescent with mutation and recombination [16]. We assume that effective population size varied according to a piecewise constant process. Following [9,10], we considered a fixed number of time windows, whose size increased exponentially from recent to old periods. More precisely, we used *I* = 21 windows of the form [*t_i_*, *t*_*i*+1_ [, where *t_i_* = exp(log(1 + *aT)i/(I —* 1)) — 1)/*a* generations BP for *i* from 0 to *I* — 1, with *T_I_* = 130, 000 and *a* = 0.06, and *t_I_* = +∞. These specific values of *T* and *a* were chosen to capture important periods of cattle history. Modifying *T* would allow population size changes to occur on a longer or shorter period in the past, and modifying a would allow to describe more precisely one specific part of the history, playing on the ratio between the length of recent versus old time windows. With our parametrization, the most recent time window ranged from present to 10 generations BP, the second most recent ranged from 10 to 25 generations BP, … the second oldest ranged from 83,000 to 130,000 generations BP and the oldest included all generations above 130,000 generations BP.

The parameters of this model are the population sizes *N_i_* for *i* from 0 to *I* — 1, the per generation per site recombination rate *r* and the per generation per site mutation rate *μ*. Prior distributions for the population sizes were taken uniform in the log 10 scale, from 10 to 100,000. In order to avoid unrealistic trajectories, we also set that the ratio of population sizes between two consecutive time windows could not exceed 10. In practice, we thus sampled log_10_(*N*_0_) uniformly between 1 and 5, and iteratively computed log_10_(N_*i*_) = max(min(log_10_(*N*_*i*-1_) + *α*, 5), 1), with *α* sampled uniformly between -1 and 1. For the recombination rate, we used an uniform prior between 1e-9 and 1e-8, consistent with recent estimations in cattle [66]. For the mutation rate we considered a fixed value, in order to compare our estimation approach with other recent ones making the same hypothesis [9,10,12], but it would be straightforward to use a prior distribution instead. This value was taken equal to 1e-8, as in [12].

### Summary statistics

We summarized each sample of *n* diploid genomes using a combination of statistics related to the allele frequency spectrum (AFS) and the average linkage disequilibrium (LD) over the genome. AFS statistics included the overall proportion of polymorphic sites over the genome (one statistic) and, among these polymorphic sites, the proportion of those with *i* copies of the minor allele, for *i* from 1 to *n* (*n* statistics). LD statistics included the average *r*^2^ over 18 different sets of SNP pairs (18 statistics), where each set was characterized by a different physical distance between SNPs. Indeed, the expected value of *r*^2^ between two SNPs at genetic distance *c* is related to the population size 1/2*c* generations BP [26]. Thus, for each of the time windows of our model, we computed *r*^2^ for SNP pairs whose physical distance would approximately correspond to a genetic distance of 1/2*t* (± 5%), where t was the middle of the window, assuming a recombination rate of 1.0 cM/Mb . For the two most recent windows, the physical distance between SNPs derived from this formula was larger than 2Mb, which could not be achieved in our simulations (see below). We thus considered only 19 statistics out of 21 windows, corresponding to distances between SNPs going from 282 bp to 1.4 Mb. We further dropped the LD statistic corresponding to a distance of 282 bp, both in the simulation study and in the real data analysis, because with our cattle data (described below) it had a strikingly low value, which was likely due to a technical problem related to the sequencing, the calling or the accuracy of the assembly. Consequently, the smallest distance bin used in our study was finally equal to 470 bp. This is specific to our study and smaller distances might be used in future studies. By default, the *r*^2^ computed between two SNPs was the zygotic LD, i.e. the correlation between the vectors of *n* genotypes observed at the two SNPs [67]. But for some comparative analyzes we also calculated the well-known gametic LD, where the correlation is computed between the two vectors of 2*n* alleles observed at the two SNPs. Note that this second option is only possible for haploid or phased data.

In many situations, we computed these summary statistics only from SNPs above a given minor allele frequency (MAF) threshold, whose value could differ between AFS and LD statistics. For a MAF threshold corresponding to c copies of the minor allele, the overall proportion of SNPs was changed to the overall proportion of SNPs with more than c copies of the minor allele, and all other proportions in the AFS were computed relative to SNPs with more than *c* copies of the minor allele. Overall, only *n* + 2 — *c* statistics were available in this case, instead of *n* + 1 without MAF threshold. In contrast, the number of LD statistics was not affected by the MAF threshold.

In a few specific analyses, we also computed summary statistics related to the distribution of IBS segment length. We summarized this distribution by a set of 11 quantiles, from 0.0001 to 1 — 0.0001.

### Implementation

We simulated 250,000 samples of 100 haploid genomes using *ms* [68], with parameters sampled from the priors described above. We chose this software because it allows simulating the exact coalescent with mutation and recombination, but faster algorithms based on approximations of this model could be used in future studies. For computational reasons, each haploid genome included only 100 independent 2Mb-long long segments. From each simulated sample of 100 haploid genomes, five different samples of *n* diploid genomes were created, for n equal to 10, 15, 20, 25 and 50. Each of these samples was created by choosing at random 2*n* haploid genomes among 100 (without replacement). In addition, 200,000 samples of 25 diploid genomes were simulated directly from *ms* samples of 50 haploid genomes. Thus, ABC analyses focusing on a sample size of 25 diploid genomes were based on 450,000 simulated samples (unless specified), while analyses involving other sample sizes were based on 250,000 simulated samples.

For the real data set, a total of 234 phased bull genomes were obtained from the 1,000 bull genomes project, Run II [40]. These included 129 Holstein (125 Black and 4 Red), 43 Fleckvieh, 47 Angus and 15 Jersey animals. Holstein animals came from various flocks with distinct geographical origins. In order to study homogeneous groups, we thus focused on the 52 Holstein animals from Australia (other geographical origins had significantly lower sample sizes). We further selected 25 unrelated animals within each breed with the following procedure: first, we removed all animals that were either extremely inbred or extremely related to another sampled animal, based on the genomic relationship matrix computed from GCTA [69]. Then, we sampled 25 animals at random among the remaining ones. For the Jersey breed, as only 15 animals were available and as they were found to be all unrelated to each other, we kept them all.

The summary statistics described above were computed using the same python script for both simulated and cattle samples. Since the length of cattle chromosomes was much larger than that of simulated segments (2Mb), we first cut each cattle chromosome into consecutive but non-overlapping 2Mb-long segments. To keep the approach computationally efficient, the average LD for a given distance bin was not evaluated from all SNP pairs satisfying the distance condition, but from a random subset of these pairs. This subset was selected by an iterative search along each 2Mb-long region, so that intervals defined by all SNP pairs did not overlap.

With the default parameter values described above, simulating 100 genomic samples and computing all summary statistics for these samples took approximately three hours on a standard computer, using a single core. Using 200 cores in parallel on a computing cluster, we could obtain 450,000 samples of summary statistics in less than 48 hours.

The final ABC estimation, based on the comparison of the simulated and observed summary statistics, was performed in R using the package *abc* [70]. By default, we accepted simulated samples with a tolerance rate of 0.005 and adjusted accepted values by a neural network regression approach [32]. This approach allows to reduce the dimension of the set of summary statistics and accounts for the non-linearity of the regression function m linking parameters and statistics (Eq. (2)). Neural network regression was applied with the default parameter values of the function *abc*, except for the final analysis of all cattle breeds where 100 (instead of 10) neural networks were fitted in order to get more stable estimations. For each parameter, a point estimate was obtained by taking the median of the posterior distribution. Variations from this default strategy were also tried, as mentioned in the results section. In particular, we also estimated posterior distributions using rejection or ridge regression [33], using the default values implemented in the *abc* package.

### Cross validation analyzes

We evaluated the performance of ABC using several subsets of summary statistics and several choices of MAF threshold, sample size, estimation approach, or tolerance. For each specific combination of these parameters, we conducted a cross validation study based on K = 2000 simulated samples, using the R function *cvabc*. The prediction error (PE) associated to a given parameter value *θ* was computed as 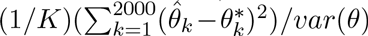 where 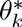 is the true value of *θ* in the *k*th simulated sample, 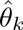is the point estimation of this value provided by ABC, and *var* (*θ*) is the prior variance of *θ*. With this scaling, estimating *θ_k_* from the prior distribution of *θ* would result in a PE of 1.

Similarly, the estimation bias for *θ* was computed as (1/K)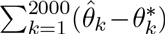, and the empirical coverage of the 90% credible interval was evaluated by (1/K)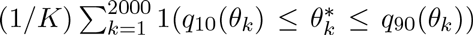, where *q*_10_(*θ_k_*) and *q*_90_(*θ*_k_) are the 5% and 95% quantiles of the posterior distribution of *θ_k_*, and 1(C) is the indicative function equal to 1 if condition *C* is satisfied and 0 otherwise.

When computing these metrics for the population size N in a given time window, we focused on parameter *θ* = log_10_(N) rather than *θ* = N. Without this rescaling, PEs and biases would only reflect the estimation accuracy for large populations, while estimation errors concerning small populations would be masked.

### Rescaling time from generations to coalescent units

Considering a population with variable population size, let *N(t)* be the haploid population size at generation *t* and 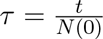 be a rescaling of time in *N*(0) units. In this time scale, the history of population size changes is summarized by the function:

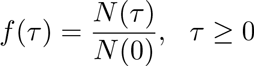

It can be shown [71] that the genealogical process of a sample of size *n* from this population, and in particular the joint distribution of all coalescence times, is identical to the genealogical process of a sample of size *n* in a constant size population where time would be rescaled by the function

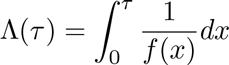

Consequently, all variable population size histories can be related to the classical Kingman’s coalescent. In this process, the expected TMRCE in a sample of size *n* is 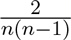 and the expected TMRCA is 2(1 — 1/n).

In S2 Fig, the PE obtained in time window [*t_i_*, *t_i_*+1 [for a given population size history was allocated to the rescaled interval [*u_i_,u_i_*_+1_[. Applying the equations above to the specific situation of a piecewise constant population size process, *u_i_* was computed as

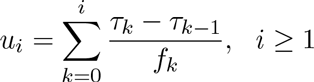

with 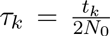 and 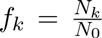 PE were then averaged over histories, for several values of u between 1e-5 and 100· Note that *N_0_* is the haploid population size here, while the population sizes mentioned anywhere else in this paper are always diploid population sizes. We used 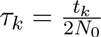, instead of the classical 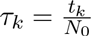 mentioned above, in order to get an expected TMRCA approximately equal to 1 (rather than 2) for large samples, which facilitates the reading of S2 Fig.

### Additional simulated datasets

Twenty scenarios with fixed population size history were considered for validation, see Fig. 3, SII Fig, S13 Fig and S14 Fig. For each of these scenarios, 20 PODs were simulated. Each of them included 25 diploid genomes and 500 independent 2Mb-long segments. Population size parameters were the same in all 20 replicates of each scenario, and the per site recombination rate was also constant and equal to 5e-9.

### Comparison of summary statistics obtained from NGS and genotyping data

For 12 of the 129 Holstein bulls considered in this study, genotypes on the 800K Illumina bovine SNP chip were obtained from the Gembal project [72]. Among the 708,771 SNPs retained in this study after quality control, 562,746 were polymorphic among the 12 bulls considered here. These SNPs were used to compute the polymorphic site AFS and the LD summary statistics from genotyping data. The rate of false negative SNPs in the NGS data was estimated by the proportion of these 562,746 positions for which no SNP was called from the NGS data. Similarly, the rate of false positive SNPs in these NGS data was estimated by considering the 145,978 SNP positions that were found monomorphic with the 800K genotypes, and computing the proportion of these positions where a SNP was called in the NGS data.

### Software and data availability

Python and R scripts for the PopSizeABC method can be found at https://forge-dga.jouy.inra.fr/projects/popsizeabc/. Simulated and observed summary statistics used in this study are also provided on this web page.

## Acknowledgments

Data analyzes were performed on the computer cluster of the Genotoul bioinformatics platform Toulouse Midi-Pyrénées (www.bioinfo.genotoul.fr). The cattle genomes were obtained from the 1,000 bull genomes project (www.1000bullgenomes.com). The 12 Holstein bull genotypes for the bovine 800K SNP chip were obtained from the GEMBAL project, which is funded by the Agence Nationale de la Recherche (ANR-10-GENM-0014), APISGENE, Races de France and INRA “AIP Bioressources”. Preliminar analyzes were performed by Stanislas Sochacki, who was funded by the Projets Exploratoires Pluridisciplinaires (PEPS 2012 Bio-Maths-Info). Flora Jay’s position was funded by the Agence Nationale de la Recherche Demochips project (ANR-12-BSV7-0012). We would like to thank Lourós Chikhi, Olivier Mazet, Simona Grusea, Bertrand Servin, Didier Boichard and all members of the Demochips project for their useful comments on this study. We are also grateful to Bertrand Servin for sharing some Python code, and to three anonymous reviewers for their constructive comments on previous versions of the manuscript.

## Supporting Information

S1 **Fig**

**Accuracy of credible intervals obtained by ABC:** Empirical coverage (left) and width (right) of the 90% credible interval for the population size in each time window. The empirical coverage is the proportion of simulated histories for which the true population size was included in the 90% credible interval of the posterior distribution. If the posterior distribution was correctly estimated, this proportion should have been 90%, as shown by the black horizontal solid line. Parameter settings were the same as in Fig. 1.

S2 **Fig**

**Accuracy of ABC estimation along the coalescent process:** Prediction error for the estimated population size when time is measured in units of the expected time to the most recent common ancestor (TMRCA) of the sample. Prediction errors were evaluated from 2,000 random population size histories. Black vertical dotted lines indicate the expected time to the most recent coalescence event, *E[TMRCE*], and the expected TMRCA, *E[TMRCA]*. Summary statistics considered in the ABC analysis were (i) the AFS and (ii) the average zygotic LD for several distance bins. These statistics were computed from *n* = 25 diploid individuals, using all SNPs for AFS statistics and SNPs with a MAF above 20% for LD statistics. The posterior distribution of each parameter was obtained by neural network regression [32], with a tolerance rate of 0.005. Population size point estimates correspond to the median of the posterior distribution.

S3 **Fig**

**Accuracy of credible intervals obtained by ABC and relative importance of the summary statistics:** Empirical coverage (left) and width (right) of the 90% credible interval for the population size in each time window. Parameter settings were the same as in Fig. 2. The very large credible intervals obtained on average with AFS statistics, in some time windows, are due to a retalively small number of PODs with extreme values.

S4 **Fig**

**Accuracy of ABC estimation based on LD summary statistics:** Prediction error for the estimated population size in each time window, evaluated from 2,000 random population size histories. Summary statistics considered in the ABC analysis were the average gametic LD (triangles) or the average zygotic LD (circles) for several distance bins. These statistics were computed from *n* = 25 diploid individuals, using different MAF thresholds. Other parameter settings were the same as in Fig. 2.

S5 **Fig**

**Influence of the number of simulated data sets on ABC estimation:** Top: Prediction error for the estimated population size in each time window (left) and standard deviation of this error (right). Bottom: Empirical coverage (left) and width (right) of the 90% credible interval for the population size in each time window. These quantiles were evaluated from 2,000 random population size histories. For each of these histories, one POD of *n* = 25 diploid genomes was simulated, where each genome consisted in 100 independent 2Mb-long segments. Population size history was estimated from this POD by ABC, for various numbers of simulated datasets (see the legend) with the same sample size (*n* = 25) and genome length (100 independent 2MB segments). Summary statistics considered in the ABC analysis were (i) the AFS and (ii) the average zygotic LD for several distance bins. AFS statistics were computed using all SNPs and LD statistics were computed using SNPs with a MAF above 20%. The posterior distribution of each parameter was obtained by neural network regression, with the tolerance rate leading to the smallest prediction error. Population size point estimates were obtained from the median of the posterior distribution.

S6 **Fig**

**Influence of the genome length of simulated and observed data sets on ABC estimation:** Top: Prediction error for the estimated population size in each time window (left) and standard deviation of this error (right). Bottom: Empirical coverage (left) and width (right) of the 90% credible interval for the population size in each time window. These quantiles were evaluated from 2,000 random population size histories. For each of these histories, one POD of *n* = 25 diploid genomes was simulated, where each genome consisted in 10, 50 or 100 independent 2Mb-long segments (see the legend). Population size history was estimated from this POD by ABC, using

450.0 simulated datasets with the same sample size (*n* = 25) and genome length. The posterior distribution of each parameter was obtained by neural network regression, with a tolerance rate of 0.005. All other settings are similar to S5 Fig.

S7 **Fig**

**Using different genome lengths for simulated and observed data sets:** Top: Prediction error for the estimated population size in each time window (left) and standard deviation of this error (right). Bottom: Empirical coverage (left) and width (right) of the 90% credible interval for the population size in each time window. These quantiles were evaluated from

2.0 random population size histories. For each of these histories, one POD of *n* = 25 diploid genomes was simulated, where each genome consisted in 10 or 100 independent 2Mb-long segments (see the legend). Population size history was estimated from this POD by ABC, using 450,000 simulated datasets with the same sample size (*n* = 25) but a possibly different genome length (see the legend). The posterior distribution of each parameter was obtained by neural network regression, with a tolerance rate of 0.005. All other settings are similar to S5 Fig.

S8 **Fig**

**Influence of the sample size on ABC estimation:** Top: Prediction error for the estimated population size in each time window (left) and standard deviation of this error (right). Bottom: Empirical coverage (left) and width (right) of the 90% credible interval for the population size in each time window. These quantiles were evaluated from 2,000 random population size histories. For each of these histories, one POD of n diploid genomes was simulated, for different values of n between 10 and 50 (see the legend). Each genome consisted in 100 independent 2Mb-long segments. Population size history was estimated from this POD by ABC, using 450,000 simulated datasets with the same sample size and genome length. All other settings are similar to S5 Fig.

S9 **Fig**

**Influence of MAF threshold on ABC estimation:** Top: Prediction error for the estimated population size in each time window (left) and standard deviation of this error (right). Middle: Bias for the estimated population size in each time window. Bottom: Empirical coverage (left) and width (right) of the 90% credible interval for the population size in each time window. These quantiles were evaluated from 2,000 random population size histories. For each of these histories, one POD of *n* = 25 diploid genomes was simulated, where each genome consisted in 100 independent 2Mb-long segments. Population size history was estimated from this POD by ABC, using 450,000 simulated datasets with the same sample size and genome length. Summary statistics considered in the ABC analysis were (i) the AFS and (ii) the average zygotic LD for several distance bins. AFS statistics were computed using different MAF thresholds, LD statistics were computed from SNPs with a MAF above 20%. The posterior distribution of each parameter was obtained by neural network regression, with a tolerance rate of 0.005. Population size point estimates were obtained from the median of the posterior distribution.

S10 **Fig**

**Estimation of population size history from the mode of the posterior distribution in six different simulated scenarios:** All settings are similar to Fig. 3, except that population size point estimates were obtained from the mode of the posterior distribution.

S11 **Fig**

**Estimation of population size history in the zigzag scenario and five related scenarios:** a scenario where all population sizes are divided by ten compared to the original zigzag (“zigzag small“, top right), a scenario where only the recent bottleneck of the original zigzag is simulated (“bottleneck1 recent large“, middle left), a scenario corresponding to the history wrongly inferred by ABC based on data from the “bottleneck1 recent large” scenario (middle right), and two scenarios where only the recent (bottom left) or the old (bottom right) bottleneck of the “zigzag small” are simulated. All settings are similar to Fig. 3.

S12 **Fig**

**Observed and best simulated summary statistics in the “bottleneck! recent large” scenario:** For one of the five PODs analyzed in this scenario, observed AFS (left) and LD (right) statistics are shown by green full circles. The average value of these statistics over the five best simulated data sets, i.e. the five simulated data sets leading to the smallest distance between observed and simulated statistics, are shown by blue crosses. The variation of these statistics over the five best simulated data sets is also indicated by blue dotted lines, which correspond to the average value plus (or minus) twice the standard deviation of each statistic.

S13 **Fig**

**Estimation of population size history in four scenarios including a bottlenck followed by a population decline:** Population size varied between 60,000 and 6,000 individuals in the top panels, and between 6,000 and 600 individuals in the bottom panels. Population size changes occurred between 2,300 and 50 generations BP in the left panels, and between 34,000 and 900 generations BP in the right panels. All settings are similar to Fig. 3.

S14 **Fig**

**Estimation of population size history in the decline scenario and five related scenarios:** a sudden (rather than continuous) decline from 40.0 to 300 individuals occurring 200 generations BP (top right), a sudden decline from 40,000 to 300 individuals occurring 1,000 generations BP (middle left), the same sudden decline followed by an expansion to 5,000 individuals occurring 580 generations BP (middle right) or an expansion to

1.0 individuals occurring 140 generations BP (bottom left), and a scenario similar to the continuous decline (top left) but including a sudden decline to 100 individuals between 230 and 140 generations BP, followed by an expansion to 1,000 individuals (bottom right). All settings are similar to Fig. 3.

S15 **Fig**

**Estimation of past effective population size using MSMC with four haplotypes in six different simulated scenarios:** For each scenario, the five PODs considered for MSMC estimation were the same as in Fig. 3. The expected TMRCA shown here is also the same as in Fig. 3, it corresponds to samples of 50 haploid sequences.

S16 **Fig**

**Estimation of past effective population size using MSMC with eight haplotypes in six different simulated scenarios:** For each scenario, the five PODs considered for MSMC estimation were the same as in Fig. 3. The expected TMRCA shown here is also the same as in Fig. 3, it corresponds to samples of 50 haploid sequences.

S17 **Fig**

**Estimation of past effective population size using MSMC with four haplotypes in the decline scenario and five related scenarios:** For each scenario, the five PODs considered for MSMC estimation were the same as in S14 Fig. The expected TMRCA shown here is also the same as in S14 Fig, it corresponds to samples of 50 haploid sequences.

S18 **Fig**

**Estimation of past effective population size using MSMC with eight haplotypes in the decline scenario and five related scenarios:** For each scenario, the five PODs considered for MSMC estimation were the same as in S14 Fig. The expected TMRCA shown here is also the same as in S14 Fig, it corresponds to samples of 50 haploid sequences.

S19 **Fig**

**Influence of phasing errors on MSMC estimation:** Estimation of past effective population size using MSMC with four haplotypes in the “small” scenario (top), the “decline” scenario (middle) and the “expansion” scenario (bottom). MSMC analyzes were run from perfectly phased data, phased data with 1 or 10 switch errors per Mb and diploid individual, or unphased data (i.e. two unphased diploid individuals). All other settings are similar to S15 Fig.

S20 **Fig**

**Comparison of summary statistics obtained from NGS and genotyping data:** polymorphic site AFS, i.e. without the overall proportion of SNPs (left), average gametic LD (middle) and average zygotic LD (right). These statistics were computed from 12 Holstein animals for which both NGS data and genotyping data were available, using only SNP positions from the 800K chip (even for the NGS data statistics). No MAF threshold was used.

S21 **Fig**

**False positive and false negative rates of SNP detection in the 1,000 bull genomes project:** Error rates were computed from 12 Holstein animals for which both NGS data and genotyping data were available. False positive SNPs were positions that were found polymorphic in the NGS data but not in the 800K data. Their minor allele count in the NGS data was called the wrong minor allele count. False negative SNPs were positions that were found polymorphic in the 800K data but not in the NGS data. Their minor allele count in the 800K data was called the true minor allele count.

S22 **Fig**

**Estimation of population size history using ABC without rare SNPs in five different simulated scenarios:** All settings are similar to Fig. 3, except that AFS statistics were computed only from SNPs with a MAF above 20%.

S23 **Fig**

**Ninety percent credible intervals of estimated population size history in four cattle breeds:** Holstein (top left), Angus (top right), Fleckvieh (bottom left) and Jersey (bottom right). Parameter settings are the same as in Fig. 6.

S24 **Fig**

**Predictive posterior check of the population size history estimated in the Holstein cattle breed (Fig. 6):** Ten thousand genomic samples were simulated under population size histories that were sampled from the posterior distribution estimated in Fig. 6. Four combinations of summary statistics were computed from each sample: AFS and LD statistics (top left), AFS statistics alone (top right), LD statistics alone (bottom left) and IBS statistics (bottom right, see the Methods for a detailed description of these statistics). For each of these combinations, a principal component analysis (PCA) of the 10,000 simulated samples was performed: the projection of all samples on the two first dimensions of this PCA are plotted in black. The vector of summary statistics observed in Holstein was then projected on the same hyperplan. It always fell within the cloud of simulated summary statistics, which shows that the estimated history is able to reproduce summary statistics that are indeed similar to the observed ones. Interestingly, this also holds for IBS statistics, which were not used for the estimation. Results are shown for the Holstein breed but they were similar for the other breeds.

S25 **Fig**

**Observed and best simulated summary statistics in the Holstein cattle breed:** Observed AFS (left) and LD (right) statistics are shown by green full circles. The average value of these statistics over the five best simulated data sets, i.e. the five simulated data sets leading to the smallest distance between observed and simulated statistics, are shown by blue crosses. The variation of these statistics over the five best simulated data sets is also indicated by blue dotted lines, which correspond to the average value plus (or minus) twice the standard deviation of each statistic.

S26 **Fig**

**Influence of population size changes on LD statistics:** LD statistics for several scenarios inplying a sudden expansion from 500 to 50,000 individuals (left) or a sudden decline from 50,000 to 500 individuals (right). Several expansion or decline times were considered, as well as two scenarios with a constant population size of 500 or 50,000 individuals (see the legend). For each scenario, LD statistics were averaged over 20 PODs including 25 diploid genomes and 100 2Mb-long regions. In contrast with expansion scenarios, some decline scenarios lead to even larger LD statistics than those obtained for a constant small population. Indeed, as these declines are very old compared to the expected TMRCA of a population of 500 individuals, their main effect is to increase, at some loci, the time during which the sample has only two ancestral lineages. Because this increase is very large (backward in time, population size, and thus expected coalescence time, are suddenly multiplied by 100), mutations occuring in this part of the coalescence tree eventually represent a large proportion of all oberved polymorphic sites. Besides, for two linked loci with similar topologies of the coalescence tree, mutations occuring in this part of the tree lead to very high *r*^2^ values, up to 1 if the topologies are exactly the same.

S27 **Fig**

**Accuracy of ABC and relative importance of LD and AFS in different families of scenarios:** Prediction error for the estimated population size in each time window, focusing on scenarios with a population size below 1.0 (top left), above 10,000 (top right), below 1,000 in the last 200 generations and above 10,000 for times more ancient than 13,000 generations BP (bottom left) or above 10,000 in the last 200 generations and below 1,000 for times more ancient than 13,000 generations BP (bottom left). For the two latter scenarios, the time window where population size goes from above 10.0 to below 1,000 (or vice versa) is delimited by vertical dotted lines. For each scenario category, PE were evaluated from 2,000 random histories. Summary statistics considered in the ABC analysis were either the AFS statistics alone, the LD statistics alone or the AFS and LD statistics together (see the legend). All other settings are similar to Fig. 2.

S28 **Fig**

**Estimation of population size history using different ABC settings in the “bottleneck1 old large” scenario:** Summary statistics considered in the ABC analysis were either the AFS statistics alone (left column), the LD statistics alone (middle column), or the AFS and LD statistics together (right column). AFS statistics were computed using either all SNPs (top panels) or only those with a MAF above 20% (bottom panels). All other settings are similar to Fig. 3.

S29 **Fig**

**Accuracy of ABC estimation based on the distribution of IBS segment lengths:** Prediction error for the population size in each time window, evaluated from 2,000 random population size histories. Summary statistics considered in the ABC analysis included several combinations of (i) the AFS, (ii) the average zygotic LD for several distance bins and (iii) the distribution of IBS segment lengths within one diploid individual. These statistics were computed from *n* = 25 diploid individuals, using all SNPs for AFS and IBS statistics and SNPs with a MAF above 20% for LD statistics. Other parameter settings are the same as in Fig. 2.

S30 **Fig**

**Added value of NGS for population size history estimation:** Estimation of population size history in the Holstein cattle breed using ABC, based on whole genome NGS data from *n* = 25 animals. Summary statistics considered in the ABC analysis included different combinations of (i) the AFS and (ii) the average zygotic LD for several distance bins. These statistics were computed either from the SNPs that are included in the 800K SNP chip or from all SNPs found in the NGS data. A MAF threshold of 20% was used for all curves and statistics. Other parameter settings are the same as in Fig. 5.

S31 **Fig**

**Population size history in Holstein using IBS statistics:** Estimation of population size history in the Holstein cattle breed using ABC, based on whole genome NGS data from *n* = 25 animals. Summary statistics considered in the ABC analysis were either both the AFS and the average zygotic LD for several distance bins, or the distribution of IBS segment lengths within one diploid individual. These statistics were computed using SNPs with a MAF above 20%. Other parameter settings are the same as in Fig. 5.

